# Fluidigm2PURC: automated processing and haplotype inference for double-barcoded PCR amplicons

**DOI:** 10.1101/242677

**Authors:** Paul D. Blischak, Maribeth Latvis, Diego F. Morales-Briones, Jens C. Johnson, Verónica S. Di Stilio, Andrea D. Wolfe, David C. Tank

**Affiliations:** Department of Evolution, Ecology, and Organismal Biology, The Ohio State University, 318 W. 12^th^ Ave., Columbus, OH, 43210-1242 USA.; Department of Natural Resource Management, South Dakota State University, 1390 College Avenue, Brookings, SD, 57007-1696 USA.; Department of Plant and Microbial Biology, University of Minnesota, 1479 Gortner Avenue, Saint Paul, MN, 55108-1095 USA.; Department of Biology, University of Washington, Seattle, WA, 98195-1800 USA.; Department of Biological Sciences, University of Idaho, 875 Perimeter Dr. MS 3051, Moscow, ID, 83844-3051 USA.; Stillinger Herbarium, University of Idaho, 875 Perimeter Dr. MS 1133, Moscow, ID, 83844-1133 USA.; Institute for Bioinformatics and Evolutionary Studies (IBEST), University of Idaho, 875 Perimeter Dr. MS 3051, Moscow, ID, 83844-3051 USA.

**Keywords:** bioinformatics, haplotype inference, high-throughput sequencing, microfluidic PCR, phylogenomics, polyploidy

## Abstract

**Premise of the study:** Targeted enrichment strategies for phylogenomic inference are a time- and cost-efficient way to collect DNA sequence data for large numbers of individuals at multiple, independent loci. Automated and reproducible processing of these data is a crucial step for researchers conducting phylogenetic studies.

**Methods and Results:** We present Fluidigm2PURC, an open source Python utility for processing paired-end Illumina data from double-barcoded PCR amplicons. In combination with the program PURC (Pipeline for Untangling Reticulate Complexes), our scripts process raw FASTQ files for analysis with PURC and use its output to infer haplotypes for diploids, polyploids, and samples with unknown ploidy. We demonstrate the use of the pipeline with an example data set from the genus *Thalictrum* L. (Ranunculaceae).

**Conclusions:** Fluidigm2PURC is freely available for Unix-like operating systems on GitHub [https://github.com/pblischak/fluidigm2purc] and for all operating systems through Docker [https://hub.docker.com/r/pblischak/fluidigm2purc].

## INTRODUCTION

The collection of large-scale, multilocus data sets for phylogenomic inference has become an increasingly common method for understanding evolutionary relationships within a group of taxa. Coupled with recent implementations of coalescent-based species tree estimation programs that take into account the independent histories of different genes (e.g., SVDquartets, Chifman and Kubatko, 2014; ASTRAL-II, Mirarab and Warnow, 2015), targeted enrichment strategies are powerful methods for collecting more informative data sets for conducting phylogenomic investigations. Of the many types of targeted enrichment that exist, several recent studies have begun to use a method that combines both library preparation and target amplification into a single step. This process, known as double-barcoded amplicon sequencing (Uribe-Convers et al., 2016), allows for the collection of multilocus sequence data for large numbers of individuals that is both time- and cost-effective.

Double-barcoded amplicon sequencing combines the amplification of a targeted region in the genome with the addition of sample-specific barcodes and Illumina sequencing adapters to the resulting PCR product for paired-end sequencing on an Illumina MiSeq platform (Uribe-Convers et al., 2016). This is done by adding conserved sequence (CS) tags to traditional PCR primers, which act as templates for adding barcodes and adapters when preparing the sequencing library. Parallel amplification is most often achieved using microfluidic PCR with the Fluidigm Access Array (Fluidigm, San Francisco, CA, USA; e.g., Gostel et al., 2015; Uribe-Convers et al., 2016; Kates et al., 2017), allowing for multiple samples and loci to be amplified simultaneously (minimum of 48 samples x 48 loci). The newer Fluidigm Juno system can also handle up to 192 samples in a single run, and multiplexing of primer pairs can allow for even higher throughput, provided that the primers do not interact during amplification. Double-barcoded amplicons can also be generated by other means using approaches such as traditional or highly-multiplexed PCR (e.g., Bybee et al., 2011; Dupuis et al., 2017).

Previous methods to analyze these data have typically relied on generating consensus sequences using software packages such as Geneious (Kearse et al., 2012; e.g. Gostel et al., 2015), HiMAP (Dupuis et al., 2017), or an R script, *reduce_amplicons.R,* that is part of the dbcAmplicons package (but see comparison with "occurrence-based" methods in dbcAmplicons in the ***Example Analyses*** section; Uribe-Convers et al., 2016; Kates et al., 2017). However, using consensus sequences can often ignore important within-individual level variation, such as differing alleles or levels of ploidy. To alleviate this issue and to facilitate the analysis of these data for haplotype inference, we developed Fluidigm2PURC. Fluidigm2PURC consists of two main Python scripts that process input data files using several external programs (Table 1) that automate quality filtering, read merging, and file formatting for downstream steps (Figure 1). Although it can be used to process any double-barcoded amplicons, the software derives its name from the method of PCR amplification that we used to generate our data (Fluidigm Access Array), as well as its primary dependency, PURC, a Python program that combines sequence clustering and PCR chimera detection (Rothfels et al., 2017). The final step in the Fluidigm2PURC pipeline processes clusters from PURC and outputs a FASTA file containing phased haplotypes for all targeted sequences. This last step has methods for haplotype inference that work on diploids, polyploids, individuals with unknown ploidy, or any mixture of the three. To demonstrate the utility of Fluidigm2PURC, we analyzed nuclear amplicon data from the genus *Thalictrum* L. (Ranunculaceae) and compared the results with those obtained from dbcAmplicons using the *reduce_amplicons.R* script (Uribe-Convers et al., 2016).

**Table 1.** Dependencies for the Fluidigm2PURC pipeline with version numbers in parentheses

| Dependency | Citation | Link |
| --- | --- | --- |
| PURC (v1.02) | Rothfels et al., 2017 | https://bitbucket.org/crothfels/purc |
| Sickle (v1.33) | Joshi and Nash, 2011 | https://github.com/najoshi/sickle |
| FLASH2 (v2.2.00) | Magoč and Salzberg, 2011 | https://github.com/dstreett/FLASH2 |
| Mafft (v7.237) | Katoh, 2013 | http://mafft.cbrc.jp/alignment/software/ |
| Phyutility (v2.7.1) | Smith and Dunn, 2008 | https://github.com/blackrim/phyutility |

**Fig. 1.**
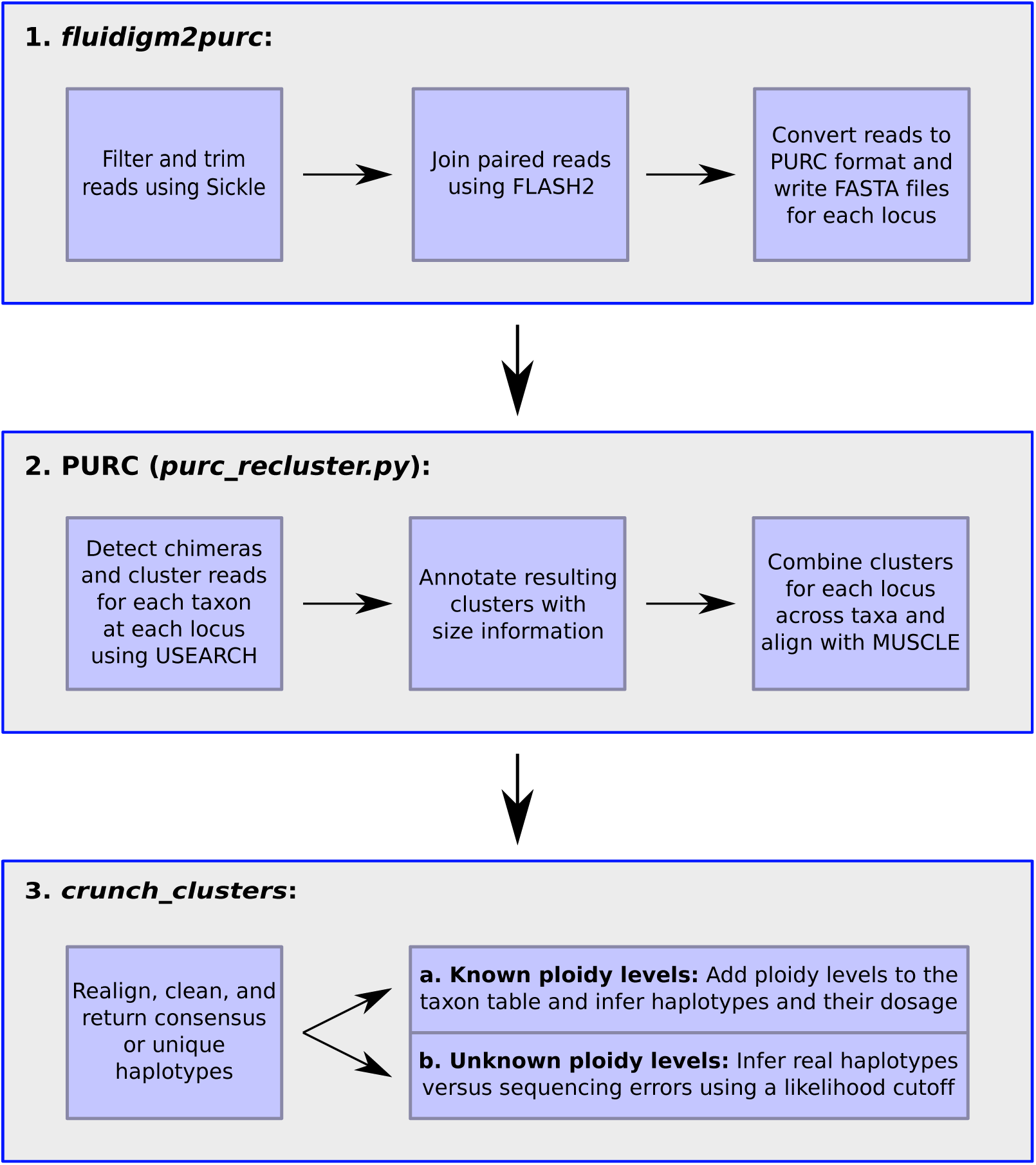
Flowchart outlining the steps for haplotype inference using Fluidigm2PURC

## METHODS AND RESULTS

***Input data***—The input data for Fluidigm2PURC are paired-end FASTQ files (R1 and R2 for paired reads) that have been demultiplexed using the program dbcAmplicons (Uribe-Convers et al., 2016). dbcAmplicons demultiplexes reads using the original sample barcodes and amplicon primer sequences to annotate the reads with the sample and locus name that each read comes from, followed by trimming these identifying parts of the sequence. The resulting pair of FASTQ files is then input into the first script in the pipeline, *fluidigm2purc*.

***Step 1: fluidigm2purc***—The *fluidigm2purc* script takes the paired-end FASTQ files, filters them using Sickle (Joshi and Nash, 2011; minimum length = 100bp, PHRED threshold = 20), merges the filtered reads using FLASH2 (Magoč and Salzberg, 2011), and then converts the resulting FASTQ files into FASTA files (one for each locus) with sequence header information that is compatible with PURC. The sequence headers for PURC follow the format ‘>IndividualName|LocusName|UniqueID#’. When paired reads with low quality bases are trimmed by Sickle and no longer overlap, we merge them artificially with multiple N’s inserted between them. The *fluidigm2purc* script writes two additional files: (1) the *taxon table*, a two-column table listing each sequenced taxon and its ploidy level, and (2) the *locus-err* table, a two-column table listing each sequenced locus and the average level of sequencing error for all reads coming from that locus. The taxon table lists the ploidy as “None” for all individuals by default, but known ploidy levels can be included by the user (e.g., diploid has the value “2,” tetraploid has the value “4,” etc.). For the locus-err table, the per locus levels of sequencing error are calculated individually from the input FASTQ files using the average PHRED score per read averaged across all reads coming from that locus.

***Step 2: PURC***—The output FASTA files from *fluidigm2purc* can be run through PURC using the *purc_recluster.py* script (Rothfels et al., 2017). This script is used to iteratively run chimera detection and sequence clustering (performed with USEARCH; Edgar, 2010; Edgar et al., 2011) on each locus individually to produce a reduced set of putative haplotypes that includes size information about the number of original reads forming each cluster. Details on running PURC can be found on its Bitbucket page [https://bitbucket.org/crothfels/purc].

***Step 3: crunch_clusters***—The clusters output by PURC are then run through our second script, *crunch_clusters*, which uses the taxon table and locus-err table output by *fluidigm2purc* (Step 1) to infer haplotypes in a maximum likelihood framework. This script also has options for realigning clusters using MAFFT (Katoh, 2013), as well as cleaning the clusters using Phyutility (Smith and Dunn, 2008).

Before haplotypes can be inferred at a locus, we first do a pairwise comparison of all clusters for each taxon individually and merge any clusters that are identical (ignoring gaps). This step is necessary because of the initial trimming/filtering step in the *fluidigm2purc* script. Artificially joining unmerged reads often causes two sequencing clusters representing the same haplotype to form: (1) one cluster for reads that were merged, and (2) one cluster for the reads that were artificially merged and contain a large number of gapped sites in the middle. In this case, these two clusters should not be treated as separate haplotypes, so we combine the clusters by keeping the larger haplotype (i.e., the one with less gaps) and adding the sizes of the two clusters together. The alternative would be to process the original data by ignoring all reads that did not merge. However, throwing away unmerged reads could potentially discard sequence variation that should be represented in the data set, especially if most reads are unmerged, which may be the case for large amplicons. The downside of merging sequences that are identical except for gaps is that it potentially discards informative indel variation, although it is unlikely that a locus within an individual would have only one of its haplotypes containing gaps and not the others. Overall, we felt that this approach provided the best method for including more of the original data when inferring haplotypes.

*Inferring haplotypes with ploidy information*—For known ploidy levels, we use a multinomial likelihood to determine the number of copies of each potential haplotype using the ordered cluster sizes returned by PURC (largest to smallest). Given an individual of ploidy level *K*, we enumerate the number of possible haplotype configurations using integer partitions (an unordered set of integers that sums to *K*; Stojmenovic and Zoghbi, 1998). Since the cluster sizes are sorted, we never need to consider more than the first *K* largest clusters. For example, a tetraploid can have a maximum of four haplotypes, and the integer partitions to consider are (4,0,0,0), (3,1,0,0), (2,2,0,0), (2,1,1,0), and (1,1,1,1). This corresponds to (4 copies of haplotype one), (3 copies of haplotype one, 1 copy of haplotype two), (2 copies of haplotype one, 2 copy of haplotype two), (2 copies of haplotype one, 1 copy of haplotype two, 1 copy of haplotype three), and (1 copy of haplotype one, 1 copy of haplotype two, 1 copy of haplotype three, 1 copy of haplotype four). The mathematical details for the likelihood function with an example calculation are presented in the Supplemental Materials (Supplemental Text §S1.1). Once the most likely configuration has been identified, the *crunch_clusters* script will return each haplotype in proportion to its representation in the maximum likelihood estimate. We have also provided options to return only unique haplotypes and to treat loci as haploid, the latter of which can be used to process organellar data. The haploid option can also be used as an alternative to finding consensus sequences for nuclear loci by returning only the cluster with the most reads.

*Inferring haplotypes without ploidy information*—For unknown ploidy levels, we no longer have information about the maximum number of haplotypes that an individual can have. However, we can use the cluster sizes to infer which clusters from PURC are actual haplotypes versus those that are likely to be sequencing errors. We do this by calculating the likelihood that each successive haplotype in the sorted list is a “real” haplotype versus a sequencing error. As an example, consider a tetraploid with six clusters identified by PURC. We first calculate the likelihood that all clusters are errors. Then we calculate the likelihood that cluster one is a real haplotype, and two through six are errors. Next, we calculate the likelihood that clusters one and two are real haplotypes, and that three through six are errors. This continues until we calculate the likelihood of all six clusters being real haplotypes. We then apply a cutoff that uses the relative increase in the likelihood when an additional haplotype is added. If treating an additional cluster as a haplotype increases the likelihood by less than the cutoff then only the previous haplotypes are kept and the others are considered errors. We use a default cutoff of 10% increase in the likelihood. An example with the likelihood function that we use for this approach is provided in the Supplemental Materials (Supplemental Text §S1.2).

***Example analysis***—To demonstrate the use of the Fluidigm2PURC pipeline, we analyzed amplicon sequence data generated from orthologs of the nuclear gene *PISTILLATA* (*PI*) in the genus *Thalictrum* L. (Ranunculaceae), which is single copy in diploids and two-copy in tetraploids (Di Stilio et al., 2005). *PI* is responsible for establishing stamen and petal identity during flower development in *Arabidopsis thaliana* (Goto and Meyerowitz, 1994), and has been used to detect reticulation in polyploid *Lepidium* L. (Brassicaceae) (Lee et al., 2002; Soza et al., 2014). Given the length of the *PI* locus, primers were designed to sequence exons 3 to 6 in two overlapping ~600-bp segments: exons 3 to 5 (*PIS_4*) and exons 4 to 6 (*PIS_3*). Our analyses focused on six species with known ploidy levels ranging from diploid (2N=2X=14) to 22-ploid (2N=22X=154). These species are presented in Table 2, with accession numbers following Soza et al. (2013). Paired-end reads were demultiplexed and annotated using dbcAmplicons (Uribe-Convers et al., 2016) followed by read trimming, merging, and sequence renaming using the *fluidigm2purc* script with default options. All reads coming from *PIS_3* and *PIS_4* were then run separately through PURC using the *purc_recluster.py* script (Rothfels et al., 2017). After clustering and chimera detection, we determined haplotypes for each amplicon using three different approaches: (1) consensus sequences using the ‘––haploid’ option, (2) unique haplotypes assuming unknown ploidy (10% likelihood cutoff), and (3) unique haplotypes using known ploidy. For each of these methods, we realigned and cleaned the sequences using MAFFT (Katoh 2013) and Phyutility (added the options ‘––realign ––clean 0.33’; Smith and Dunn, 2008).

**Table 2.** *Thalictrum* L. species included in the comparison of Fluidigm2PURC and dbcAmplicons. Collection information is listed as the collector(s), collection number, and the herbarium. All additional information is available from Soza et al. (2013), Tables S1, S3, and S4. For all analyses, *T*. *pubsescens*was analyzed at the 22X level.

| Species | Ploidy Level | Collection Information |
| --- | --- | --- |
| *T. thalictroides* (L.) A.J. Eames & B. Boivin | 2N=2X=14 | V. Di Stilio, 123, WTU |
| *T. squarrosum* Stephan ex Willd. | 2N=6X=42 | V. Di Stilio & X. Duan *Thalictrum* sp#8, 20120617, PE |
| *T. macrostylum* Shuttlew. ex Small & A. Heller | 2N=8X=56 | R. Penny (unvouchered) |
| *T. pubescens* Pursh | 2N=12X=84 or<br>2N=22X=154 | D. Baum & D. Howarth, 375, A |
| *T. revolutum* DC. | 2N=20X=140 | V. Soza, 1917, WTU |
| *T. dasycarpum* Fisch., C. A. Mey. & Avé-Lall. | 2N=22X=154 | V. Di Stilio, 110, WTU |

As a comparison, we also analyzed these data using the *reduce_amplicons.R* script from the dbcAmplicons package (v0.8.5; Uribe-Convers et al., 2016). This script merges paired-end reads using FLASH2 (Magoč and Salzberg, 2011) and allows for a global read trimming size to be used for read one, read two, or both. Unmerged reads are treated independently, resulting in separate haplotypes for read one and read two. The final result is a FASTA file with the unaligned haplotypes that can be further processed for downstream applications. We generated consensus haplotypes as well as haplotypes based on read occurrence (controlled by the minimum read frequency and minimum read count) using the default settings, and trimmed 20 bp from read one and 40 bp from read two. We then aligned the resulting sequences using Mafft (Katoh, 2013). These results were compared to the haplotypes from Fluidigm2PURC based on (1) the number of recovered haplotypes, (2) the length of the resulting alignment, and (3) the amount of gaps in the alignment.

*Results*—Haplotypes inferred by both methods were visualized and compared using alignment statistics computed in Geneious v8.1.8 (Kearse et al., 2012) and MEGA v7.0.18 (Kumar et al., 2016). Consensus sequences from the Fluidigm2PURC and dbcAmplicons pipelines were similar overall, with the *reduce_amplicons.R* script producing longer haplotypes, but containing more gaps (Table 3). We then compared the occurrence-based method from the *reduce_amplicons.R* script with the *crunch_clusters* results when ploidy levels are treated as unknown. In this case, Fluidigm2PURC recovered more haplotypes with fewer gaps and more parsimony informative sites. We believe the reason that the *reduce_amplicons.R* script recovered so few haplotypes is due to its use of minimum read count and frequency criteria that rely on reads being identical to form haplotypes, rather than clustering based on similarity. Inferring haplotypes with Fluidigm2PURC using known ploidy levels resulted in the largest number of recovered haplotypes. The reason that using known versus unknown ploidy levels produced more haplotypes (*PIS_3*: 57 vs. 18, *PIS_4*: 43 vs. 14) was because the clusters sizes that went into the likelihood calculation were disparate for some species (a few large clusters and many smaller ones), making the smaller clusters difficult to model when the ploidy level was unknown due to lack of prior knowledge about how many haplotypes should be expected. On a per species basis, using known ploidy levels always led to more inferred haplotypes (Table 4). For example, the *PIS_4* region for *Thalictrum pubescens* recovered 15 haplotypes when assuming known ploidy (analyzed as 22X), but only one haplotype when assuming unknown ploidy. The reason for this is that the cluster data for this species had one putative haplotype with many reads (147), but all other putative haplotypes had far fewer reads (the next largest cluster had 25 reads, and nine clusters had fewer than 10 reads). In general, drawing the line between real haplotypes and errors for clusters with lower read counts is difficult when the ploidy level is unknown. By applying a threshold, the method we implement is a conservative way to estimate haplotypes that only includes clusters with the highest read counts.

**Table 3.** Overall alignment statistics for the comparison between Fluidigm2PURC and the *reduce_amplicons.R* script.

|  | *PIS_3* |  | *PIS_4* |  |
| --- | --- | --- | --- | --- |
|  | Fluidigm2PURC | *reduce_amplicons* | Fluidigm2PURC | *reduce_amplicons* |
| *Consensus* |  |  |  |  |
| Alignment length (bp) | 395 | 415 | 403 | 418 |
| Percent gaps | 27.9 | 30.0 | 1.9 | 4.4 |
| Parsimony informative sites | 19 | 19 | 11 | 10 |
| *Unknown Ploidy/Occurrence* |  |  |  |  |
| Number of haplotypes | 18 | 5 | 14 | 6 |
| Alignment length (bp) | 395 | 403 | 403 | 424 |
| Percent gaps | 16.2 | 48.9 | 2.7 | 7.1 |
| Parsimony informative sites | 48 | 11 | 43 | 10 |
| *Known ploidy* |  |  |  |  |
| Number of haplotypes | 57 | – | 43 | – |
| Alignment length (bp) | 395 | – | 403 | – |
| Percent gaps | 10.3 | – | 2.6 | – |
| Parsimony informative sites | 81 | – | 62 | – |

**Table 4.** Per species data for the number of haplotypes inferred by Fluidigm2PURC using known vs. unknown ploidy. Data are presented as: number of inferred haplotypes (average percent gaps per haplotype). For *Thalictrum pubescens*, haplotypes are presented at both the 12X and 22X level.

| Species | Ploidy | *PIS_3* |  | *PIS_4* |  |
| --- | --- | --- | --- | --- | --- |
|  |  | Known | Unknown | Known | Unknown |
| *T. thalictroides* | 2N=2X=14 | 1 (2.3) | 1 (2.3) | 1 (0.8) | 1 (0.8) |
| *T. squarrosum* | 2N=6X=42 | 6 (3.3) | 4 (3.1) | 4 (3.9) | 3 (3.2) |
| *T. macrostylum* | 2N=8X=56 | 8 (13.9) | 5 (20.9) | 4 (3.3) | 4 (3.3) |
| *T. pubescens* | 2N=12X=84 or | 12X: 11 (3.7) | 1 (1.5) | 12X: 9 (2.3) | 1 (1.2) |
|  | 2N=22X=154 | 22X: 19 (3.3) |  | 22X: 15 (2.3) |  |
| *T. revolutum* | 2N=20X=140 | 10 (26.9) | 4 (40.9) | 5 (2.5) | 3 (2.9) |
| *T. dasycarpum* | 2N=22X=154 | 13 (9.2) | 3 (2.8) | 14 (2.4) | 2 (2.1) |

*Code and data availability*—The code for each step of our example analysis is available in the Supplemental Materials (Supplemental Text §S2). Raw sequence data from the *PIS_3* and *PIS_4* loci for the six sampled *Thalictrum* species, as well as all output FASTA files from the Fluidigm2PURC and dbcAmplicons pipelines, are available on Dryad (dryad.####).

## CONCLUSIONS

The ability to infer haplotypes regardless of an individual’s ploidy level is a crucial step toward understanding the complex relationships within many plant groups whose evolutionary histories often contain multiple instances of hybridization and whole genome duplication (Soltis and Soltis, 2009; Van de Peer et al., 2009). As models that accommodate these processes continue to be developed (e.g., Jones et al., 2013; Solís-Lemus and Ané, 2016; Oberprieler et al., 2017; Thomas et al., 2017; Wen and Nakhleh, in press), we anticipate that the functionality of our pipeline will be especially useful for conducting phylogenomic studies with nuclear sequence data. Furthermore, the increase in genomic resources for taxa across the Plant Tree of Life will continue to facilitate the process of phylogenetic marker development, allowing more researchers to take advantage of targeted enrichment strategies such as double-barcoded amplicon sequencing. Compared with existing approaches for analyzing these data, the methods we present here offer an improved workflow for sequence processing, clustering, and haplotype inference, and are particularly well suited for analyses in taxa with incomplete knowledge about ploidy levels.

***Availability***—Fluidigm2PURC is open source software that is freely available on GitHub [https://github.com/pblischak/fluidigm2purc] for Unix-like operating systems (Mac, Linux) under the GNU General Public License v3. We have also built a Docker image with all dependencies (Table 1) pre-installed for use on any operating system with a compatible distribution of the Docker software [https://hub.docker.com/r/pblischak/fluidigm2purc] (https://www.docker.com; Merkel, 2014). Fluidigm2PURC is written in Python and has been successfully tested using Python versions 2.7 and 3.6. Documentation for the software can be found on ReadTheDocs [http://fluidigm2purc.readthedocs.io].

## ACKNOWLEDGEMENTS

The authors thank C. Rothfels for helpful discussions regarding the use of PURC. We also thank S. Uribe-Convers for providing valuable feedback while testing the Fluidigm2PURC code. This work was supported by the following grants from the National Science Foundation (NSF): DEB-1455399 (ADW, L. S. Kubatko), DEB-1253463 (DCT), and IOS-1121669 (VSD), with additional support to DCT and VSD from the NSF BEACON Center for the Study of Evolution in Action (DBI-0939454).

